# A bifunctional coiled-coil protein generates the membrane-within-condensate architecture of the CO_2_-fixing pyrenoid

**DOI:** 10.64898/2026.06.09.731149

**Authors:** Sabrina L. Ergun, Claire Dignazio, Jonathan Bouvette, Haoyu Wu, Eric Franklin, Claire D. McWhite, Martin C. Jonikas

## Abstract

How membranes are integrated into biomolecular condensates is a fundamental question in cell biology. In the algal pyrenoid, an organelle responsible for one-third of global carbon fixation, CO_2_-delivering thylakoid membranes must penetrate a phase-separated condensate of the CO_2_-fixing enzyme Rubisco, but the mechanism governing membrane recruitment into the condensate remains unknown. Here, we demonstrate that the *Chlamydomonas reinhardtii* protein MITH1 acts as a molecular anchor that brings membrane into the pyrenoid condensate. MITH1 dimerizes into an extended coiled coil with an N-terminal amphipathic helix that binds thylakoid membrane. The coiled coil contains multiple novel binding sites for the Rubisco large subunit, allowing the condensate to wet onto the membrane. The coiled coil extends away from the membrane and promotes membrane organization within the condensate. These findings solve a longstanding mechanistic question in pyrenoid biogenesis and reveal general principles for how membranes are integrated into biological condensates.

## Main Text

Biological condensates and membranes are the two fundamental organizers of cellular space, yet the principles governing their functional integration remain poorly understood. While it is increasingly recognized that membrane surfaces can template condensate assembly^1–4^ and, conversely, that condensates can exert mechanical force to remodel membranes^5–9^, the mechanisms by which lipid bilayers are recruited into and organized within a biological phase-separated condensate are largely unknown.

The algal pyrenoid, a specialized CO_2_-fixing organelle responsible for approximately one-third of global carbon assimilation^10–13^, offers a unique window into this problem. Its function depends on its unusual architecture, which consists of a phase-separated condensate^14^ traversed by membranes^15,16^ (Fig. 1A). The pyrenoid’s phase-separated condensate consists primarily of Rubisco, the enzyme that fixes CO_2_ into biomass^17^, while the specialized traversing membranes release concentrated CO_2_ to enhance Rubisco’s activity^18,19^. The membranes are essential to pyrenoid function^19^, likely because they provide a compartmentalized, low-pH space that promotes CO_2_ release from bicarbonate^18,20,21^. Despite the essential importance of the membrane-within-condensate architecture for pyrenoid function, the molecular basis for how membranes are brought into the Rubisco condensate remains unknown.

**Fig. 1.**
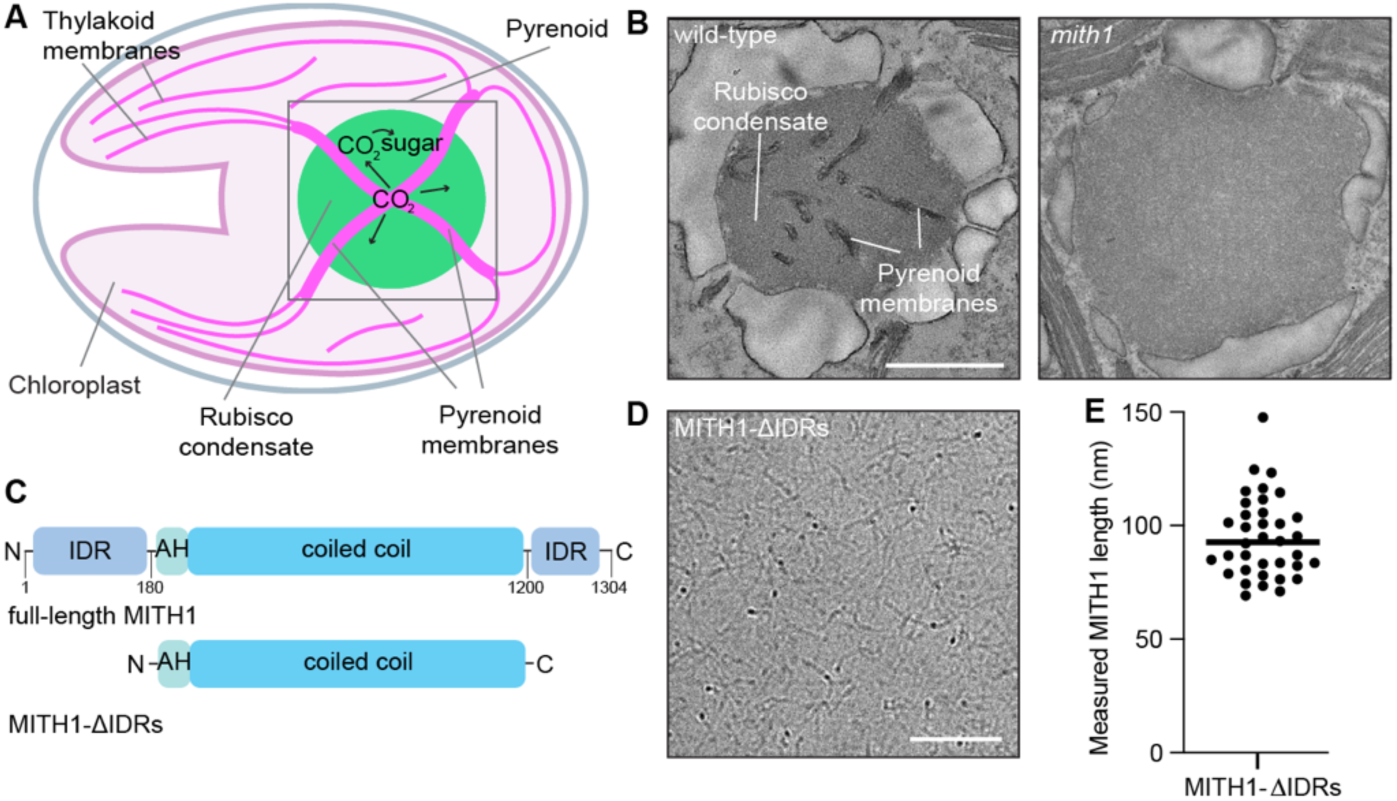
| MITH1 forms an extended coiled coil. **A** Cartoon depicting the Chlamydomonas pyrenoid, composed of the Rubisco condensate and the membrane tubules. CO_2_ is provided to Rubisco via the membrane tubules, which are essential for pyrenoid function. **B** Transmission electron micrograph (TEM) of a wild-type Chlamydomonas pyrenoid and a *mith1* mutant pyrenoid. Scale bar is 1 µm. **C** Diagram of the domains of full-length MITH1 and MITH1-ΔIDRs. **D** Cryogenic-electron microscopy (cryo-EM) micrograph of MITH1-ΔIDRs. Scale bar is 100 nm. **E** Quantification of the length of the MITH1 fibers in the cryo-EM images.

Previously, we found that the pyrenoid protein MITH1 (Cre06.g259100) is required for the formation of pyrenoid-traversing membranes^19^ in the leading model alga *Chlamydomonas reinhardtii*^22^ (Chlamydomonas hereafter). In wild-type Chlamydomonas, pyrenoid-traversing membranes form tubules throughout the condensate. In the absence of MITH1, the Rubisco condensate still forms, but pyrenoids lack a normal tubule network (Fig. 1B), suggesting that MITH1 plays a direct role in pyrenoid-traversing membrane biogenesis^19^. However, MITH1 lacks transmembrane domains and known pyrenoid condensate-interaction motifs^23^. Thus, the mechanism of MITH1’s contribution to pyrenoid-traversing membrane biogenesis has remained a central puzzle in pyrenoid biology.

Here we demonstrate that MITH1 brings membranes into the pyrenoid condensate through a unique wetting activity mediated by an amphipathic helix, an extended coiled-coil domain, and a non-canonical Rubisco-binding motif, suggesting general principles for how membranes can be brought into and organized within biomolecular condensates.

## Results

### MITH1 is an extended coiled-coil protein

Sequence analysis previously identified predicted coiled-coil regions along the length of MITH1, flanked by disordered regions at both ends^24^ (Fig. 1C). MITH1’s coiled-coil regions could fold back on themselves, or they could form an extended coiled coil. To distinguish between these possibilities, we obtained cryo-electron micrographs of purified MITH1 lacking its disordered termini (MITH1-ΔIDRs, Fig. 1C, Supplementary Fig. 1A). We observed that MITH1-ΔIDRs forms extended filament-like structures (Fig. 1D) with an average length of approximately 93 nm (Fig. 1E), which roughly corresponds to the predicted length of a fully extended coiled coil of MITH1-ΔIDRs (∼110 nm)^25^. These results suggest that MITH1 forms an extended coiled coil with disordered regions at both ends.

### MITH1 binds membrane via an amphipathic helix

Having established the overall geometry of MITH1, we then sought to determine how each region of the protein supports its function. Our previous work demonstrated that MITH1 localizes to the pyrenoid tubules^19^, but the molecular basis for this interaction was unknown. To determine if MITH1 can bind membranes directly, we used an *in vitro* liposome flotation assay, where a sucrose gradient is used to separate liposomes and associated proteins from soluble species (Fig. 2A)^26^. Purified MITH1-ΔIDRs was enriched in the top layer of the liposome flotation assay in comparison to BSA, a control soluble protein, indicating that MITH1 can bind membranes directly (Fig. 2B, C).

**Fig. 2.**
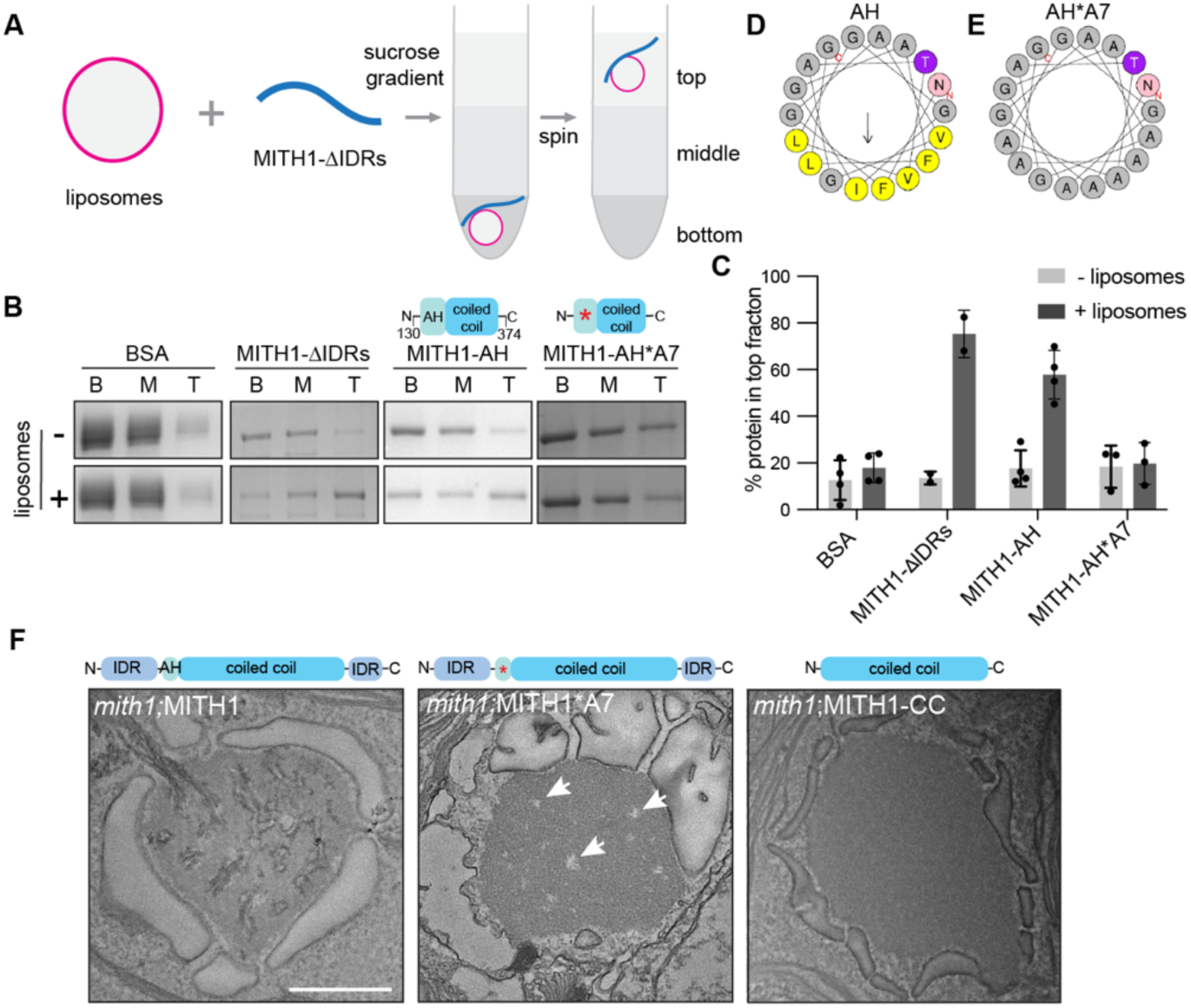
| MITH1 binds membrane via an amphipathic helix. **A** Cartoon depicting a liposome flotation assay. Liposomes and purified proteins are mixed and layered at the bottom of a sucrose gradient. The gradient is spun, liposomes float to the top, and enrichment in the top fraction indicates protein-liposome interaction. **B** SDS-PAGE of the bottom, middle, and top fractions of the flotation assay. **C** Bar graph showing the enrichment of proteins in the liposome-containing fraction relative to the bottom fraction of the flotation assay of several replicates. Error bars indicate s.d. **D** Helical wheel depicting the amphipathic helix near the MITH1 N terminus. **E** Helical wheel containing residues mutated in MITH1-AH*A7. **F** TEMs of pyrenoids expressing the indicated protein variant. Arrows indicate gaps in condensate in the *mith1;MITH1*A7* strain. Scale bar is 1 µm.

MITH1’s direct binding to membrane was unexpected because previous work had not revealed any obvious membrane-binding regions. Motivated by the above findings, we performed a bioinformatic search for potential membrane-binding motifs (Supplementary Fig. 1B)^27^. The search identified a hydrophobic sequence at the N-terminus of MITH1’s coiled-coil domain containing a hydrophobic moment, suggesting it could act as an amphipathic helix (Fig. 2D, Supplementary Fig. 1B). Amphipathic helices mediate contacts with membranes^28^, and several notable long coiled-coil proteins mediate their functions through amphipathic helix-based membrane contacts, such as the Golgi vesicle tethers Golgins and several of the mitochondrial structural Mic proteins^29,30^. The predicted amphipathic helix of MITH1 does not contain many charged residues on the non-membrane side of the helix, unlike several well-characterized amphipathic helices whose charged residues interact with lipid head groups in the membrane^31^. However, ∼90% of thylakoid lipid head groups are uncharged^32^ and therefore charged residues may not be required for thylakoid membrane binding.

To determine if MITH1’s predicted amphipathic helix mediates membrane binding, we purified a 180-amino acid segment of MITH1 containing this helix (MITH1-AH, Supplementary Fig. 1C). MITH1-AH promoted the formation of membrane tubules *in vitro* (Supplementary Fig. 1E), a classic indication of amphipathic helix-mediated membrane binding^33^. MITH1-AH bound membranes directly in our liposome flotation assay (Fig. 2B, C), while mutating the seven hydrophobic residues within the amphipathic helix to alanine (MITH1-AH*A7, Fig. 2E and Supplementary Fig. 1D) abolished membrane binding (Fig. 2B, C). We thus conclude that MITH1 binds directly to membranes via its N-terminal amphipathic helix.

To determine whether the binding of MITH1 to membrane via its amphipathic helix is important for its cellular function, we sought to determine whether the same hydrophobic residues impact the formation of the tubule network in vivo. While expressing full-length wild-type MITH1 rescued the tubule network of the *mith1* mutant^19^ (Fig. 2F), expressing a full-length MITH1 with mutated hydrophobic residues (MITH1*A7) did not rescue tubule networks (Fig. 2F), despite the observation that the MITH1*A7 protein was expressed and localized to the pyrenoid (Supplementary Fig. 2A). Together, these data indicate that the direct binding of MITH1 to membrane via its amphipathic helix is necessary for pyrenoid tubule biogenesis.

### MITH1 clears space in the condensate

While the MITH1*A7-expressing strain lacked pyrenoid-traversing membranes, it exhibited small Rubisco-depleted gaps in the condensate reminiscent of pyrenoid tubule patterning (Fig. 2F, indicated by arrows, Supplementary Fig. 3C), that were not observed in the *mith1* mutant strain (Fig. 1B, Supplementary Fig. 3B). The observation of these gaps in the condensate, together with the result that MITH1*A7 retained a non-homogeneous localization pattern as observed for wild-type MITH1-Venus (Supplementary Fig. 2A, C), suggests that even in the absence of membrane binding, MITH1 can self-organize within the condensate and contribute to clearing space in the condensate for membrane entry. In a strain expressing only the MITH1 coiled-coil domain (MITH1-CC, Supplementary Fig. 2B), gaps in the Rubisco condensate were not observed (Fig. 2F, Supplementary Fig. 3D), suggesting that the MITH1 IDRs may be necessary for this clearing function.

### The MITH1 coiled coil promotes tubule penetration into the condensate

Having established a role of membrane binding for the N-terminus of MITH1, we then sought to determine the role of the long coiled-coil domain in tubule formation. To this end, we generated a MITH1 construct lacking the coiled-coil domain (MITH1-ΔCC, Fig. 3A, D) and expressed it in a *mith1* mutant strain. The construct was expressed at a significantly lower level than that of full-length MITH1 (roughly one-tenth of the total fluorescent signal above background), did not rescue strain growth of the *mith1* mutant, and localized to one to two puncta adjacent to the pyrenoid (Fig. 3E, Supplementary Figs. 2D and 4). TEMs showed that this strain completely lacked membranes in the pyrenoid, indicating that the long coiled-coil domain of MITH1 is essential for MITH1’s stability and function (Fig. 3E, F, Supplementary Fig. 3A, E).

**Fig. 3.**
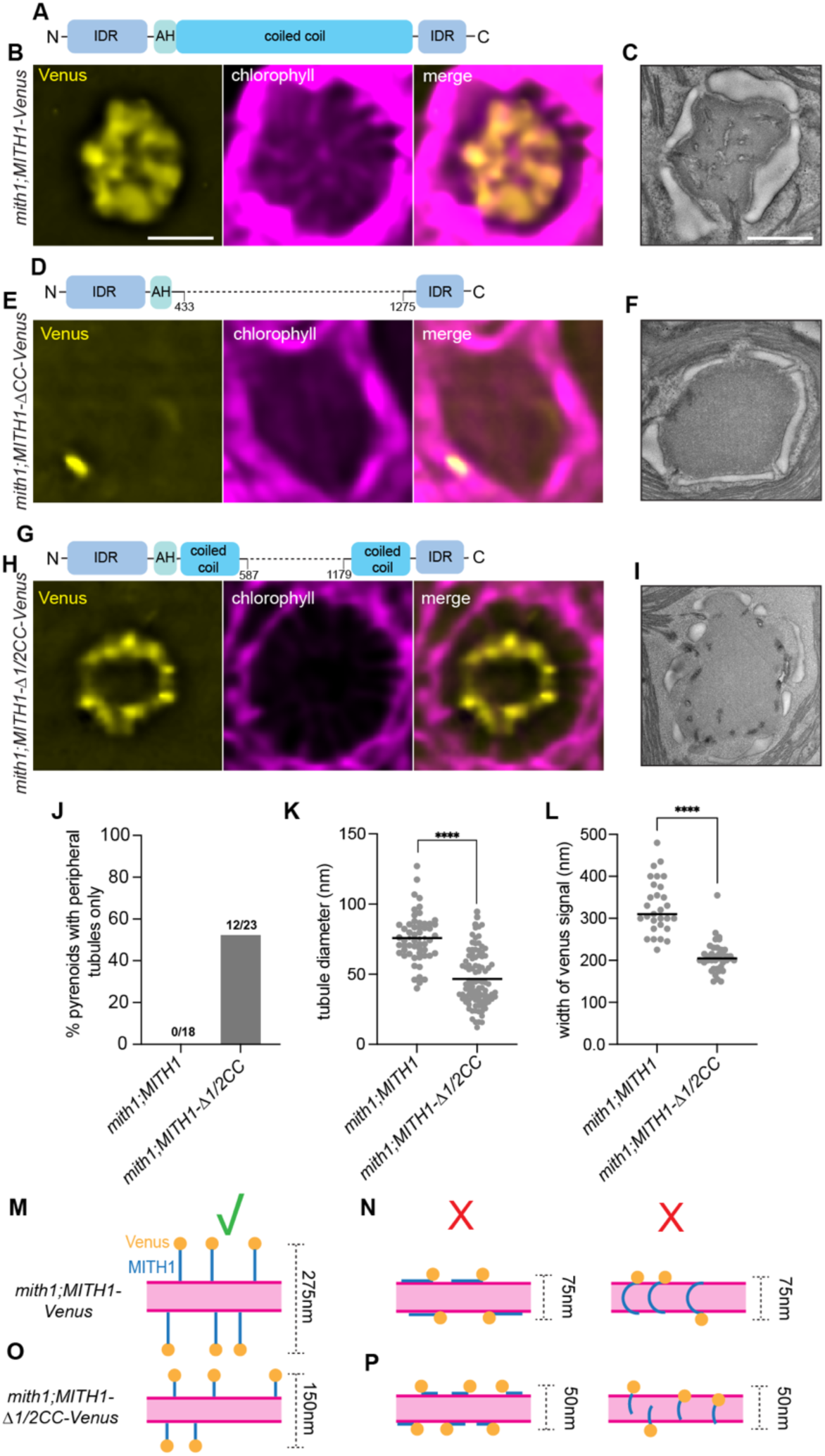
| MITH1’s coiled coil extends away from the membrane and promotes tubule penetration into the condensate. **A-I** TEM and confocal microscopy of the pyrenoids of strains expressing Venus-tagged MITH1 variants in the *mith1* mutant background. A diagram of the MITH1 variant is depicted above each set of images. Scale bars are 1 µm. **J** Quantification of the percentage of pyrenoids containing only peripheral tubules. N is shown above each bar. **K** Quantification of the tubule diameter in the indicated strains. 60-90 individual tubules were measured from 10-12 cells. **L** Quantification of the Venus signal width in the indicated strains. 35-50 individual tubules were measured from 8-10 cells. **M-P** Model of how the MITH1 coiled-coil domain could be oriented with respect to the tubule membrane and the expected Venus signal width in each case. The tubule is in magenta, MITH1 is depicted in blue, and Venus is depicted as an orange ball at the C terminus of MITH1. **** = p-value < 0.0001.

**Fig. 4.**
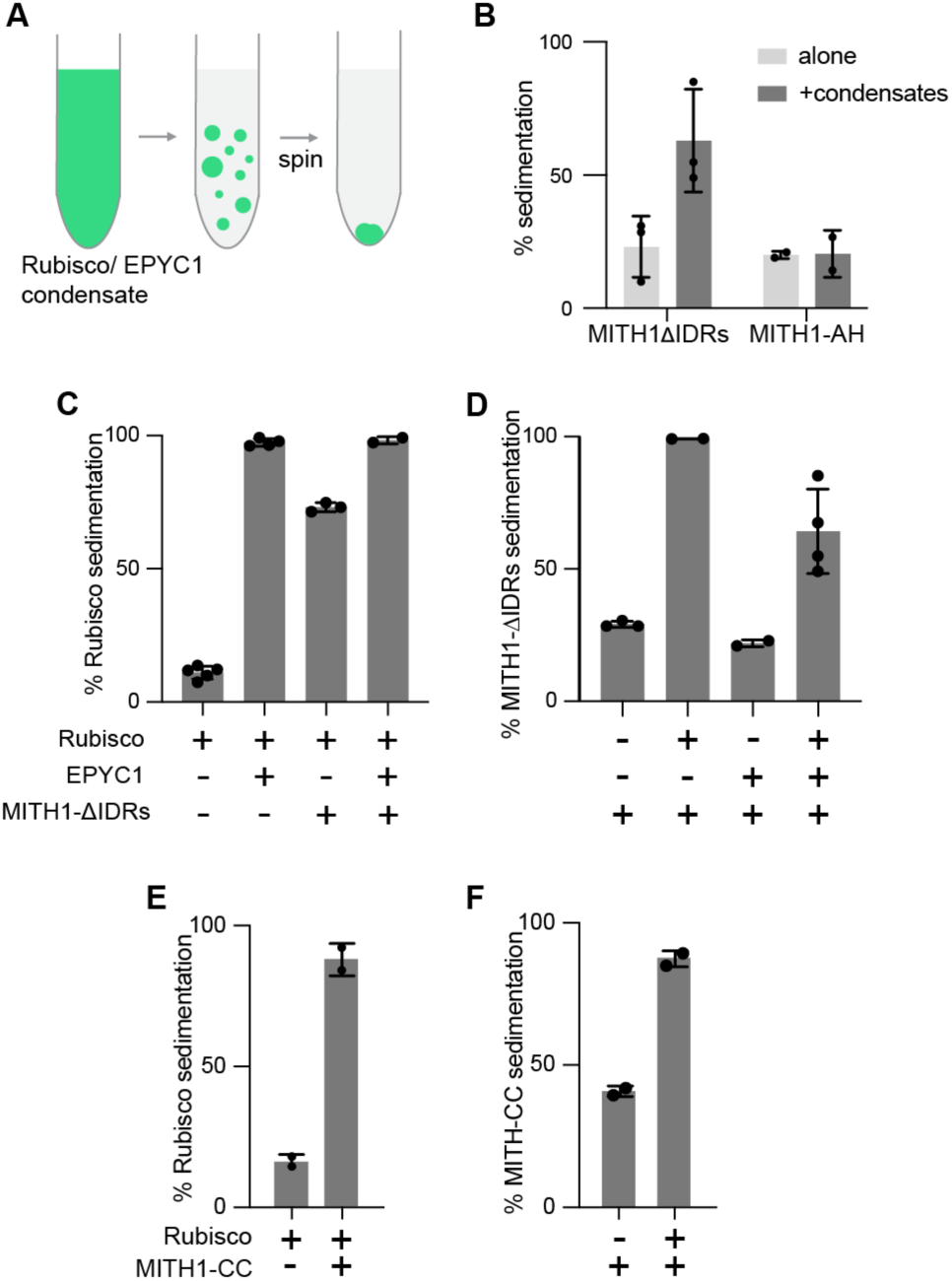
| The MITH1 coiled-coil domain co-sediments with Rubisco. **A** Diagram of an *in vitro* sedimentation assay. When spun, proteins that form a condensate will sediment to the bottom of the tube. **B** Bar graph of the percent of each protein that sedimented to the pellet when combined with the Rubisco/EPYC1 condensate *in vitro*. **C, D** Bar graphs of the sedimentation of MITH1-ΔIDRs and Rubisco in the presence or absence of MITH1-ΔIDRs, Rubisco, or EPYC1. **E, F** Same as **(C and D)** with the MITH1 coiled-coil domain only.

Due to the difficulty of obtaining high expression of the MITH1-ΔCC construct, we sought to interrogate the function of the MITH1 coiled-coil domain by truncating it. We therefore generated a strain expressing MITH1 lacking the central half (400 amino acids) of the coiled coil (*mith1;MITH1-Δ1/2CC*, Fig. 3G). This construct expressed at levels similar to those of full-length MITH1, rescued growth of the *mith1* mutant strain at air and very low CO_2_ levels, and showed pyrenoid tubule localization (Fig. 3H, Supplementary Figs. 4 and 2E). However, in 12 of 23 TEM sections of pyrenoids from this strain (∼52%), the tubules were absent from the central region of the pyrenoid (Fig. 3I, J, Supplementary Fig. 3F), a phenotype we never observed in cells expressing the full-length MITH1 construct (0/18 pyrenoids, 0%). These results suggest that the full length of the MITH1 coiled-coil domain is necessary for robust tubule penetration into the condensate.

We additionally observed that the tubules in the MITH1-Δ1/2CC strain were narrower than those within the full-length MITH1-expressing strain. When quantified, we observed a 25 nm (∼30%) decrease in average tubule diameter between the MITH1 and the MITH1-Δ1/2CC strains (Fig. 3K), suggesting that the length of the coiled-coil domain of MITH1 affects tubule diameter. Overall, these data support a role for the MITH1 coiled-coil domain in promoting normal tubule diameter and membrane penetration into the condensate.

### MITH1 extends away from the membrane into the condensate

Our structural data indicate that MITH1 is an elongated coiled coil anchored to the membrane by its N-terminus. This geometry allows for two distinct orientations: the coiled coil could extend radially away from the membrane into the condensate (Fig. 3M), or it could lie along the membrane surface (Fig. 3N). We had previously noted that the fluorescent streaks of Venus signal of full-length MITH1-Venus (Fig. 3B, Supplementary Fig. 2C) were substantially broader than those of other Venus-tagged tubule membrane proteins (such as RBMP1-Venus) or autofluorescence from tubule-associated chlorophyll^19,23^. In light of our new structural data, this observation supports an orientation where MITH1’s coiled coil extends radially away from the membrane (Fig. 3M), as the coiled coil would then provide a space between the C-terminal Venus signal and the membrane. Further supporting MITH1 extending radially away from the membrane, the MITH1-Δ1/2CC-Venus strain containing a shortened coiled-coil domain displays a reduced width of the Venus fluorescence signal (Fig. 3H, L, Supplementary Fig. 2E). If MITH1 were to lie flat on the membrane, the fluorescence width would be defined by the membrane tubule width regardless of the protein length (Fig. 3N, P). The observed signal widths also quantitatively match a model of radial extension. The full-length MITH1 signal width is consistent with a ∼75 nm tubule flanked by ∼100 nm filaments (total ∼275 nm), while the MITH1-Δ1/2CC signal width fits a ∼50 nm tubule flanked by ∼50 nm filaments (total ∼150 nm) (Fig. 3M-P). These data demonstrate that the MITH1 coiled coil extends away from the membrane to engage the condensate.

### The MITH1 coiled-coil domain co-sediments with Rubisco

Since our data suggested that MITH1 extends into the condensate, we sought to test if MITH1 directly interacts with condensate components. To test for an interaction of MITH1 with the Rubisco condensate, we utilized a co-sedimentation assay^34^ (Fig. 4A). The two main components of the pyrenoid condensate are Rubisco and its linker protein EPYC1^35^, which phase-separate *in vitro* at known concentrations^36^. Using this assay, we showed that purified MITH1-ΔIDRs can co-sediment with the Rubisco/EPYC1 condensate (Fig. 4B), demonstrating that the MITH1 coiled-coil domain interacts with at least one of the pyrenoid condensate components.

To determine if MITH1 interacts with Rubisco, EPYC1, or both, we measured co-sedimentation in pairwise interactions. MITH1 and Rubisco alone co-sedimented in the absence of EPYC1 (Fig. 4C, D), indicating that MITH1 and Rubisco can directly interact. By contrast, MITH1 and EPYC1 alone did not co-sediment (Fig. 4C, D). Together, these data indicate that MITH1 physically interacts with Rubisco but not EPYC1.

The 180-amino acid segment of MITH1 containing the amphipathic helix (MITH1-AH) did not co-sediment with Rubisco and EPYC1 (Fig. 4B), suggesting that MITH1’s coiled-coil domain is responsible for the interaction with Rubisco. Indeed, we found that just the MITH1 coiled-coil domain co-sedimented with Rubisco (Fig. 4E, F, Supplementary Fig. 5A), indicating that the extended coiled-coil domain of MITH1 contains Rubisco binding site(s).

### MITH1 binds the Rubisco large subunit

To identify the Rubisco binding site(s) on MITH1, we tested Rubisco binding to a peptide array of 25-amino acid peptides tiling across the MITH1 sequence (Fig. 5A, Supplementary Fig. 5B)^37^. We observed binding to nine sites along the coiled-coil domain, all containing a common motif [L/I]XXXXDL[K/R]QXLNXX[K/R] (Fig. 5A, B). We also observed binding to two sites within the N-terminal IDR of MITH1 consisting of a positively charged patches containing 4–7 lysine or arginine residues. To validate these MITH1 binding sites, we measured the binding of peptides representing them to Rubisco using fluorescence anisotropy^38^. A 15-amino-acid peptide containing the common motif bound Rubisco robustly and showed dose-dependent binding with an affinity in the low micromolar range, compared to no interaction with control protein BSA (Fig. 5C). In contrast, peptides representing the two positively charged patches from the N-terminus of MITH1 did not show any binding to Rubisco by fluorescence anisotropy (Fig. 5D), suggesting that the observed Rubisco binding to these peptides on the array may have been an artifact of the local enrichment of positive charges due to the crowded nature of the peptides on the array. We conclude that MITH1 can directly bind Rubisco at multiple sites along its coiled-coil domain that each harbor a common interaction motif.

**Fig. 5.**
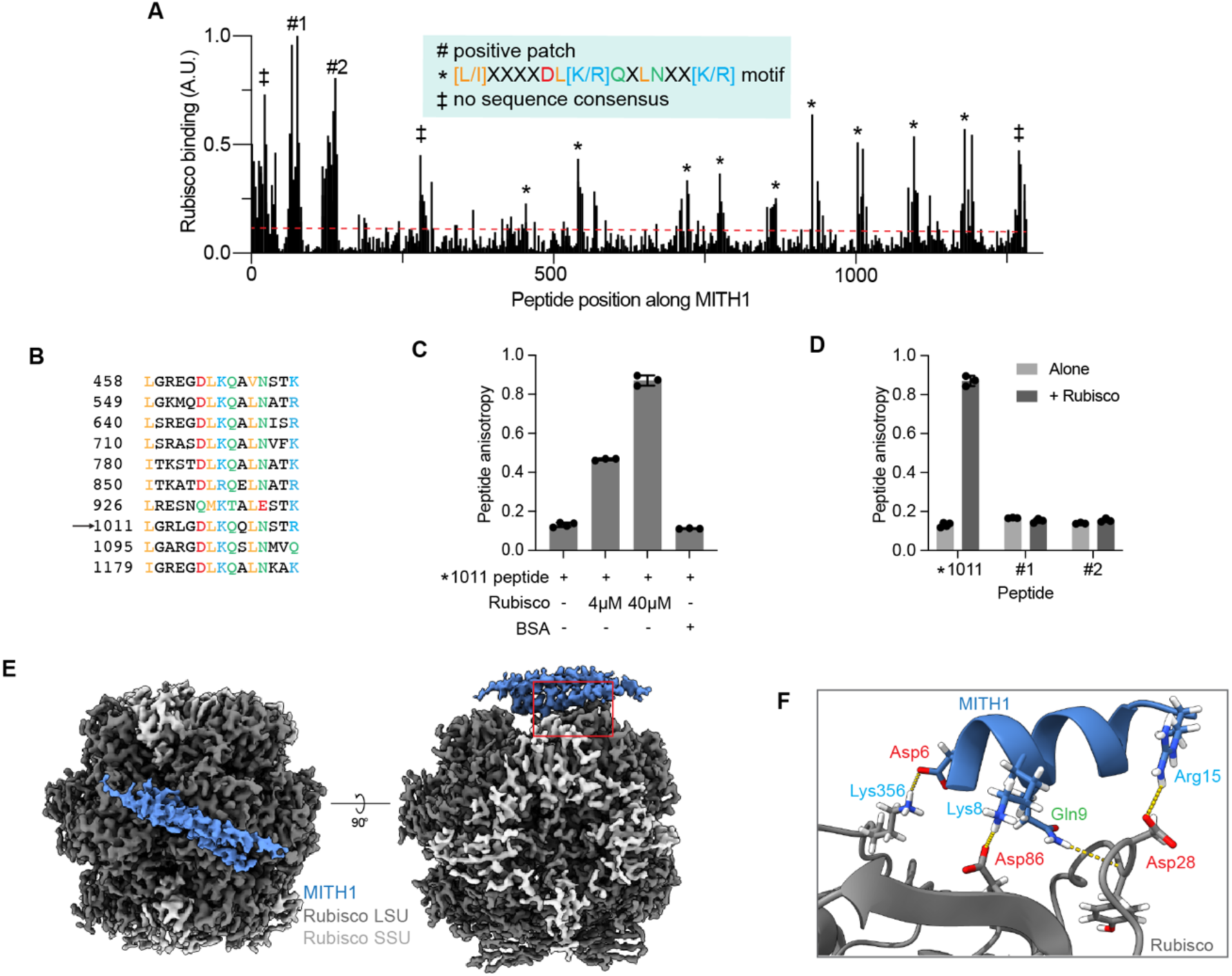
| MITH1 binds the Rubisco large subunit. **A** Rubisco-binding signal as a function of peptide position along MITH1, as measured by a peptide tiling array. The dashed red line represents the signal obtained from negative control peptides. The sequence consensus between binding sites is indicated. **B** Alignment of motif-containing Rubisco binding repeat sequences along MITH1. The arrow indicates the peptide used for anisotropy measurements. **C, D** Binding of peptides containing either the Rubisco-binding consensus sequence on MITH1 or the two N-terminal positive patches to Rubisco or BSA control measured by fluorescence anisotropy. **E** Cryo-EM map of Rubisco with the MITH1 coiled-coil domain bound to the large subunit. **F** Residues involved in the MITH1:Rubisco interaction. Error bars depict s.d.

To further elucidate how MITH1 binds Rubisco, we solved a high-resolution single-particle cryo-electron microscopy structure of Rubisco in complex with MITH1-ΔIDRs (Fig. 5E). While Rubisco was fully resolved in the structure, we could only resolve the region of MITH1 in close proximity to Rubisco, presumably due to the flexibility of MITH1. The structure shows that MITH1 binds the Rubisco large subunit, adjacent to the subunit’s catalytic site^39^. This binding site is novel and distinct from the previously known site where all other established Rubisco-binding pyrenoid factors such as EPYC1, SAGA1, RBMP1, and RBMP2 bind, which is located on the Rubisco small subunit^23,40^. This distinct binding site may allow MITH1 to diffuse through the Rubisco condensate without directly competing with other Rubisco-binding pyrenoid factors.

In the structure, a clear density consisting of two MITH1 helices is visible, demonstrating that MITH1-ΔIDRs forms a coiled-coil dimer. We could only resolve the MITH1 peptide to ∼4 Å, whereas we resolved Rubisco to ∼2.5 Å, in part due to partial occupancy of binding sites on Rubisco and the averaging across the different MITH1 binding sequences. We therefore used the sequence of the validated MITH1 Rubisco-binding peptide (Fig. 5B) to model the MITH1 residues. On the Rubisco large subunit, Asp28, Asp86, and Lys356 formed salt bridges with an arginine, lysine, and aspartic acid, respectively, within the binding motif on MITH1 (Fig. 5F). Taken together, the data demonstrate that MITH1 binds directly to the Rubisco large subunit, via multiple Rubisco-binding sites along the MITH1 coiled-coil domain.

MITH1 binding to the Rubisco large subunit has implications for the ability to engineer a pyrenoid into land plants, a recent focus of the field with the critical goal of improving crop yield^18,41–43^. In recent efforts, pyrenoid condensate-traversing membranes were reconstituted in *Arabidopsis thaliana* by expressing MITH1 together with other factors, but the reconstitution was inefficient, with only 32% of pyrenoid condensates in the best plant lines containing traversing membranes^19^. Following our discovery that MITH1 directly binds to Rubisco, we observed that the *Arabidopsis* Rubisco large subunit lacks a critical MITH1-binding-site residue (D86H) and is unable to bind MITH1 (Supplementary Fig. 5C). This incompatibility potentially explains the inefficient reconstitution of pyrenoid-traversing membranes in *Arabidopsis* and provides a path for overcoming it by engineering improved Rubisco-MITH1 binding.

### MITH1 wets condensate onto membrane *in vitro*

Thus far we have shown that MITH1 can directly interact with the thylakoid membrane via its amphipathic helix, and with the Rubisco condensate via multivalent interactions on its coiled-coil domain. These observations led us to hypothesize that MITH1 could act as a molecular anchor that promotes adhesion between the membrane and the condensate within the pyrenoid. To test this hypothesis, we designed an *in vitro* assay centered around the concept of wetting, the biophysical property of droplets spreading along a surface due to favorable binding interactions^44^. In our assay, we added MITH1-ΔIDRs to a mixture of liposomes and Rubisco/EPYC1 condensates to determine whether MITH1 can promote wetting (Fig. 6A). Upon addition of MITH1-ΔIDRs, we observed a clear wetting of the condensates on the liposome surface, whereas without MITH1-ΔIDRs the condensates only peripherally associated with the liposomes, possibly due to weak electrostatic interactions (Fig. 6B, Supplementary Fig. 6). We quantified the wetting in two ways. First, we found that addition of MITH1-ΔIDRs caused a loss of circularity of the condensates from 0.93 to 0.74 (p<0.0001, unpaired t-test, two-tailed) as they spread along the membrane (Fig. 6C). Second, upon addition of MITH1-ΔIDRs we observed a decrease in the contact angle between condensate and membrane from 118°±17° to 69°±20° (mean ± s.d., p<0.0001, unpaired t-test, two-tailed), again indicating the wetting of the condensate along the membrane (Fig. 6D)^45^. These data establish that MITH1 is a bifunctional protein anchor that mediates adhesion between the thylakoid membrane and the Rubisco condensate.

**Fig. 6.**
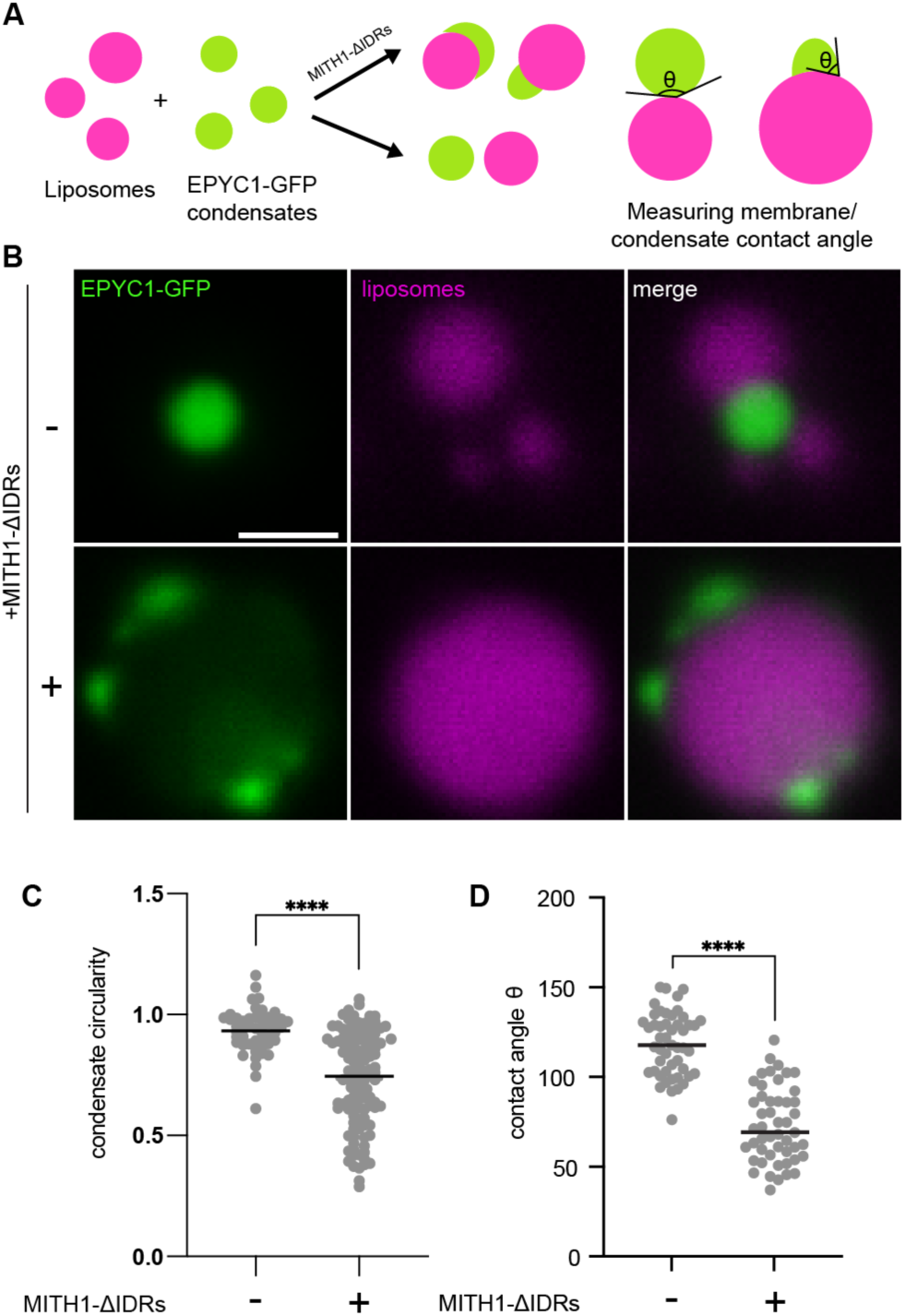
| MITH1 wets condensate onto membrane *in vitro.* **A** Overview of the condensate and membrane interaction assay. Liposomes and EPYC1-GFP/Rubisco condensates were mixed and imaged by confocal microscopy with and without MITH1-ΔIDRs, and the contact angle between the condensate and membrane was measured. **B** Confocal microscopy images of the liposomes and condensates together with and without purified MITH1-ΔIDRs. Scale bars are 1 µm. **C** Quantification of the circularity of condensates in both conditions. n ≥ 60. Unpaired t-test, two-tailed. **D** Quantification of contact angle between the condensate and the membrane. n ≥ 30. Unpaired t-test, two-tailed. **** = p value < 0.0001.

## Discussion

Here we identify MITH1 as the molecular anchor that couples thylakoid membranes to the Rubisco condensate, explaining how the pyrenoid achieves its defining membrane-within-condensate architecture. We show that MITH1 forms an extended dimeric coiled-coil that tethers the thylakoid membrane to the Rubisco condensate: its N-terminal amphipathic helix binds to the membrane, while its coiled-coil domain projects into the condensate to engage multiple Rubisco holoenzymes via salt bridge interactions with the large subunit (Fig. 7A). This bifunctional tethering allows MITH1 to wet the condensate onto the membrane surface, driving the formation of the traversing membrane network essential for efficient CO_2_ fixation.

**Fig. 7.**
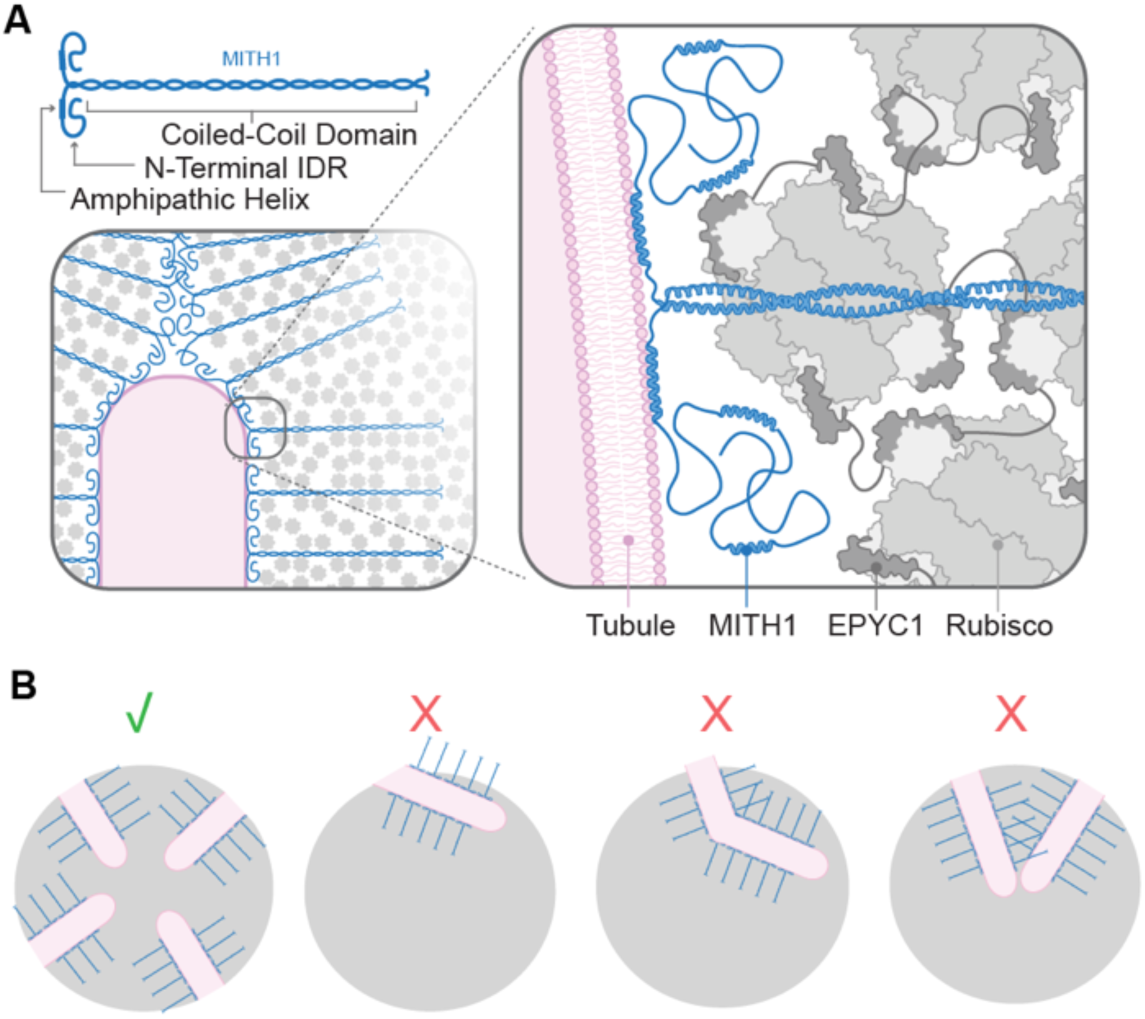
| Proposed mechanism for how MITH1 generates the membrane-within-condensate architecture of the pyrenoid. **A** Model of the structural role of MITH1 as a molecular anchor promoting tubule and condensate interaction in the pyrenoid. **B** Illustrations of how MITH1’s architecture could promote radial tubule entry into the pyrenoid and inter-tubule spacing, and disfavor tubule bending.

Our observations suggest that MITH1’s IDRs create gaps in the condensate that may promote membrane entry. The MITH1*A7 amphipathic helix mutant retained MITH1’s non-homogeneous localization within the condensate (Supplementary Fig. 2A) and displayed gaps within the condensate that, while lacking membranes, were of similar diameter to and spacing as the pattern produced by membrane tubules in the wild-type condensate (Fig. 2F). This observation suggests that the MITH1 in this mutant still self-organizes within the condensate into a configuration similar to that achieved when tubules are present. The dependence of the observed condensate gaps on the MITH1 IDRs (Fig. 2F) and the orientation of MITH1 with its N-terminus towards tubules (Fig. 3M) together suggest that MITH1’s N-terminal IDR generates the gaps in the Rubisco condensate. Indeed, IDRs have been previously shown to have the capacity to exclude proteins from condensates^46^. We speculate that the clusters of MITH1’s N-terminal IDR could locally exclude Rubisco, creating a less protein-dense region that facilitates membrane entry. The clusters of MITH1 N-termini would represent “hotspots” of membrane-binding amphipathic helices that would provide high membrane affinity to promote membrane entry into the condensate. Upon membrane binding by the amphipathic helix, the MITH1 N-terminal IDRs would fold back toward the condensate, potentially contributing to the Rubisco-free zone observed in close proximity to the membranes^47^. An amphipathic helix allows for a reversible membrane-binding interaction^48^ that we speculate may be essential to enable rapid pyrenoid dynamics and self-organization during cell division^14^.

Our data show that beyond enabling membrane entry, MITH1 organizes membranes within the condensate. We found that the coiled-coil domain of MITH1 was necessary for normal membrane organization within the condensate, as a shortened MITH1 coiled coil impaired full tubule penetration into the center of the pyrenoid (Fig. 3H-J). MITH1’s role in membrane organization can be explained by our observation that MITH1’s semi-rigid coiled coil extends away from the membrane (Fig. 3M). MITH1’s extension away from the membrane could promote radial entry of tubules into the condensate, as tubule entry at any other angle would result in unoccupied Rubisco binding sites on MITH1 protruding from the condensate (Fig. 7B). The propensity of MITH1 to extend away from the membrane tubules may also promote their continuation radially into the condensate by disfavoring bending of the tubules, as a bent tubule would lead to increased steric clashes between adjacent MITH1 molecules on the same tubule (Fig. 7B). Additionally, the propensity of MITH1 to extend away from tubules may enforce a minimum distance between adjacent tubules via steric clash (Fig. 7B). Future characterization of the three-dimensional morphology of the tubule network in MITH1 truncation mutants could test how MITH1’s coiled coil promotes tubule radial entry, controls tubule persistence length, and acts as a “ruler” that sets internal pyrenoid length scales.

By defining the molecular mechanism by which MITH1 promotes entry of membranes into the *Chlamydomonas* pyrenoid, we provide a paradigm for this critical step in the biogenesis of an organelle responsible for approximately one-third of global carbon fixation. Beyond this fundamental insight, the identification and structural resolution of the MITH1:Rubisco interface provides a blueprint for overcoming a longstanding barrier to engineering a pyrenoid into plants to enhance crop yields. On a broader cellular scale, our discovery that a bifunctional extended coiled-coil protein can mediate wetting, promote entry of membranes into a condensate, and direct their internal organization contributes conceptual insights and a model system for further study of the underlying principles of this class of cellular structures.

## Supplementary Figures

**Fig. S1.**
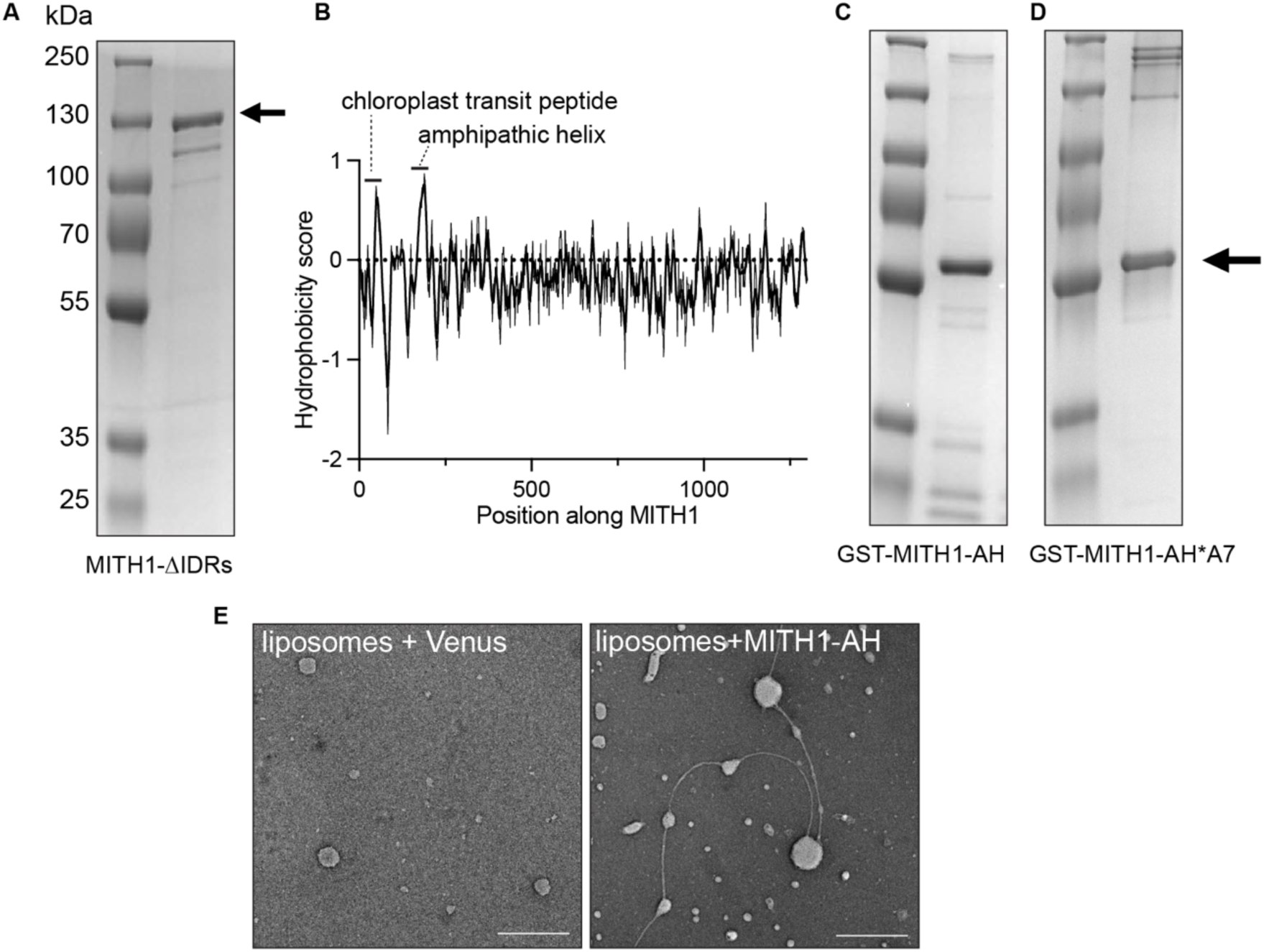
| MITH1 is a long coiled-coil protein that binds membrane via an amphipathic helix. **A** SDS-PAGE showing purified MITH1-ΔIDRs. **B** Measurement of the hydrophobicity and the hydrophobic moment along the length of MITH1. The location of the chloroplast transit peptide and the site of the amphipathic helix are indicated. **C** SDS-PAGE of the purified MITH1-AH-containing segment of MITH1. **D** SDS-PAGE of the MITH1-AH with 7 hydrophobic residues of its amphipathic helix mutated to alanine (MITH1-AH*A7). **E** Negative stain TEMs of liposomes with and without the addition of the MITH1-AH segment or Venus as a negative control. The scale bar is 500 nm.

**Fig. S2.**
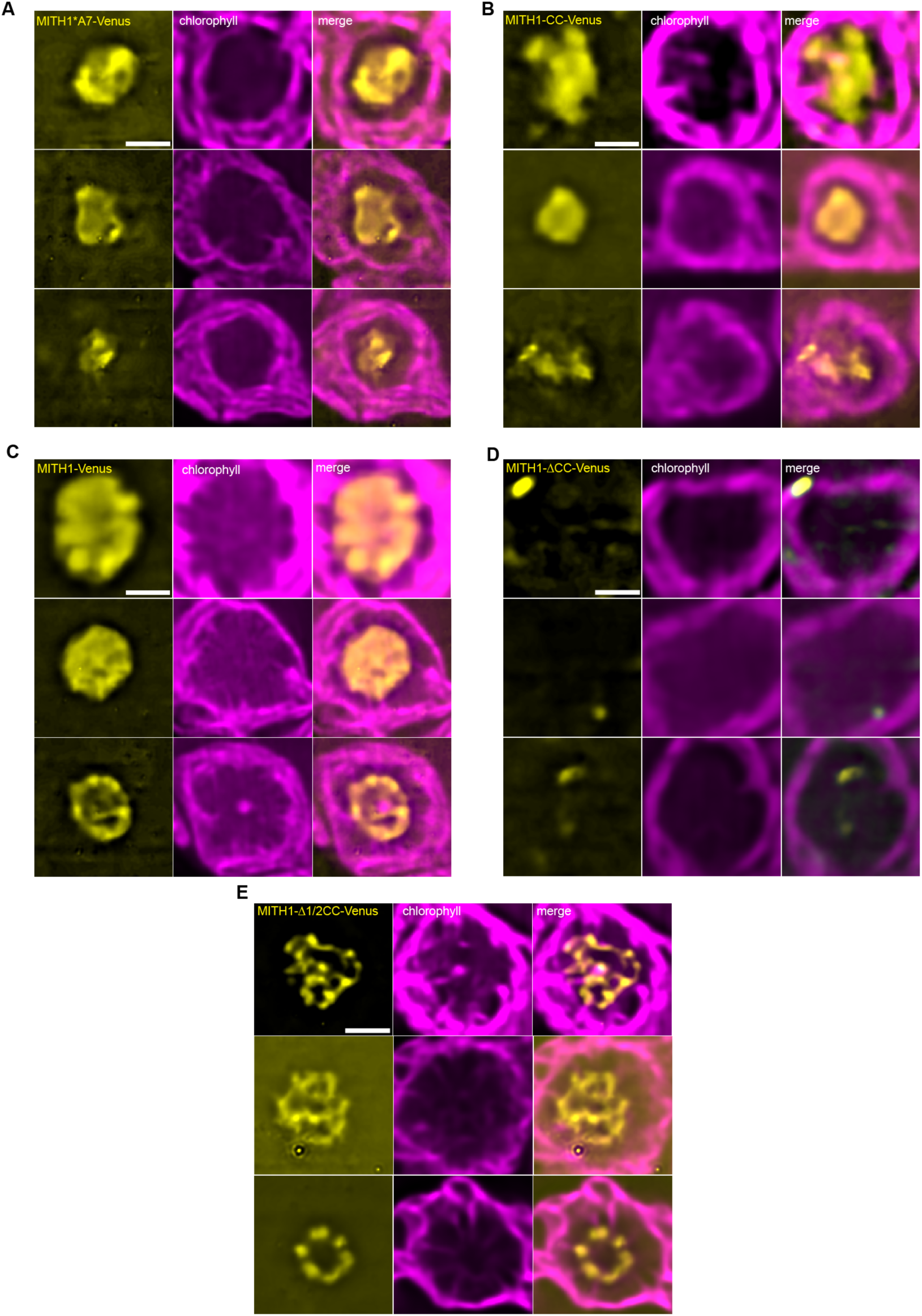
| Confocal microscopy images of MITH1 strains. A-E. Confocal microscopy of *mith1* Chlamydomonas pyrenoids expressing the indicated MITH1 variants. Scale bars are 1 µm.

**Fig. S3.**
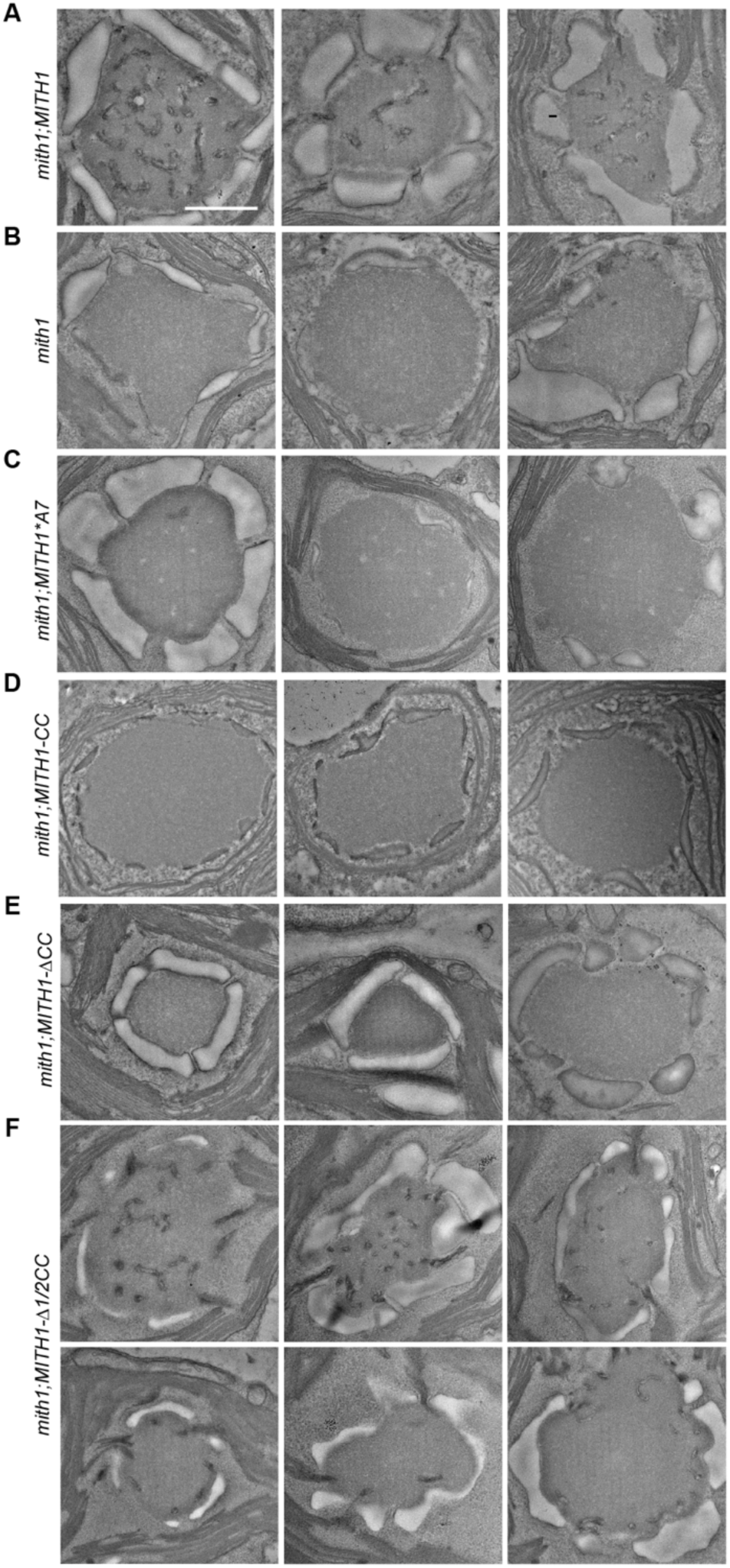
| Transmission electron microscopy of MITH1 strains. A-F. Additional examples of TEMs of pyrenoids expressing the indicated MITH1 variants. Scale bar is 1 µm.

**Fig. S4.**
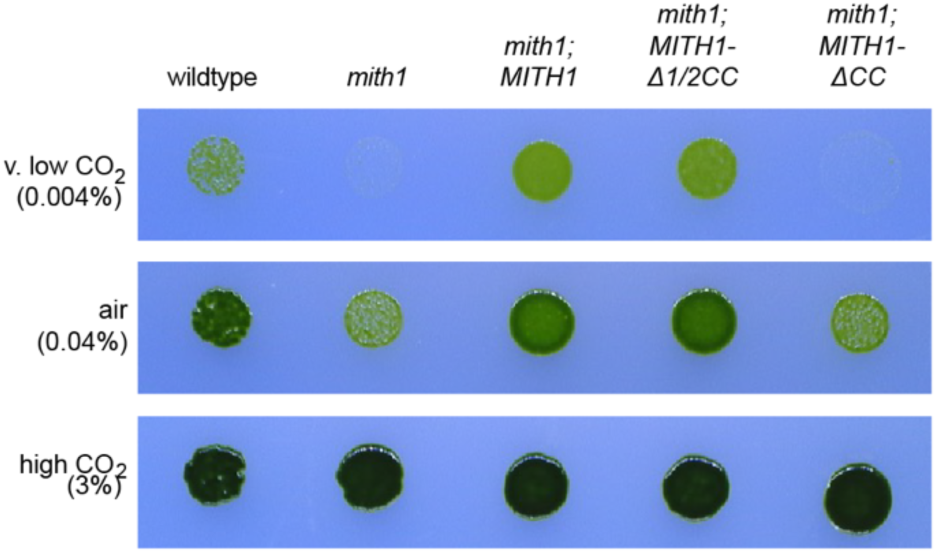
| Spot test growth assay of Chlamydomonas strains expressing the indicated MITH1 truncations.

**Fig. S5.**
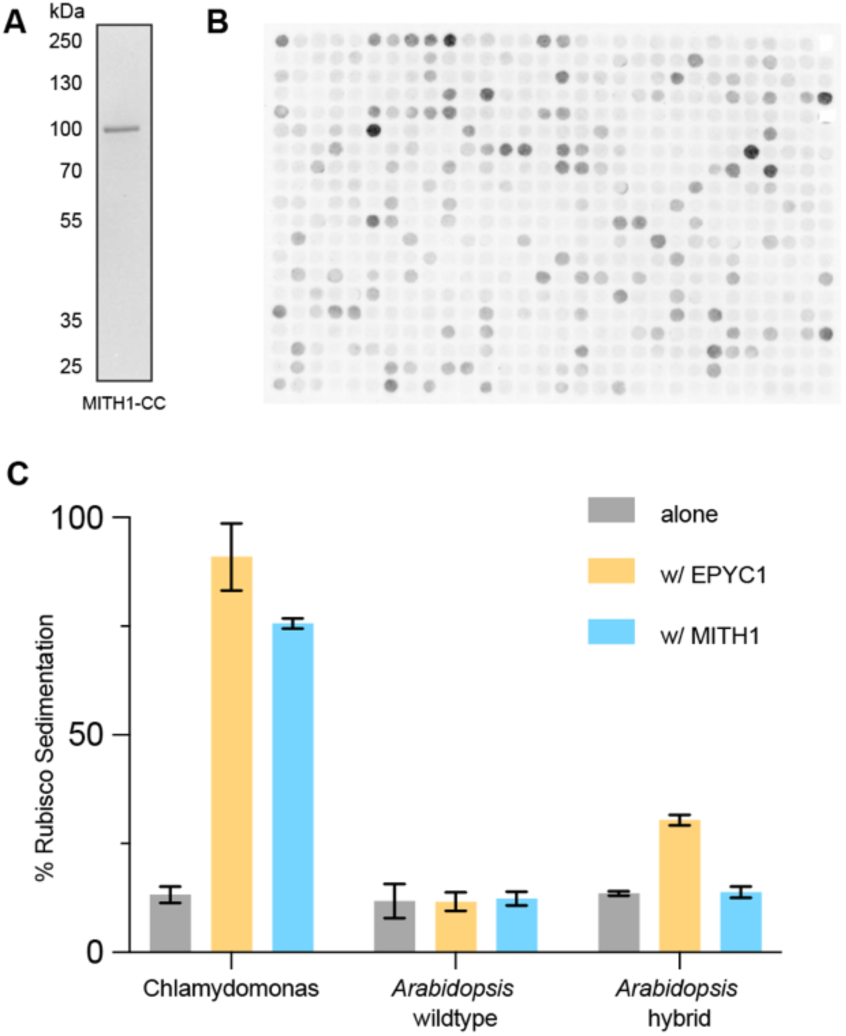
| MITH1 coiled-coil domain binds Rubisco large subunit. **A** SDS-PAGE of the MITH1-CC construct purified from Sf9 insect cells. **B** Raw image of Rubisco bound to the peptide binding array. Each spot contains many copies of one 25-amino-acid MITH1 peptide. The peptides are tiled every three amino acids along MITH1 and their position on the membrane is randomized. Every fourth peptide is repeated twice. **C** Sedimentation assay of MITH1-ΔIDRs or EPYC1 with Rubisco from Chlamydomonas, native Arabidopsis (Col-0), or Arabidopsis hybrid expressing the Chlamydomonas small Rubisco subunit (CrRbcS2)^49^.

**Fig. S6.**
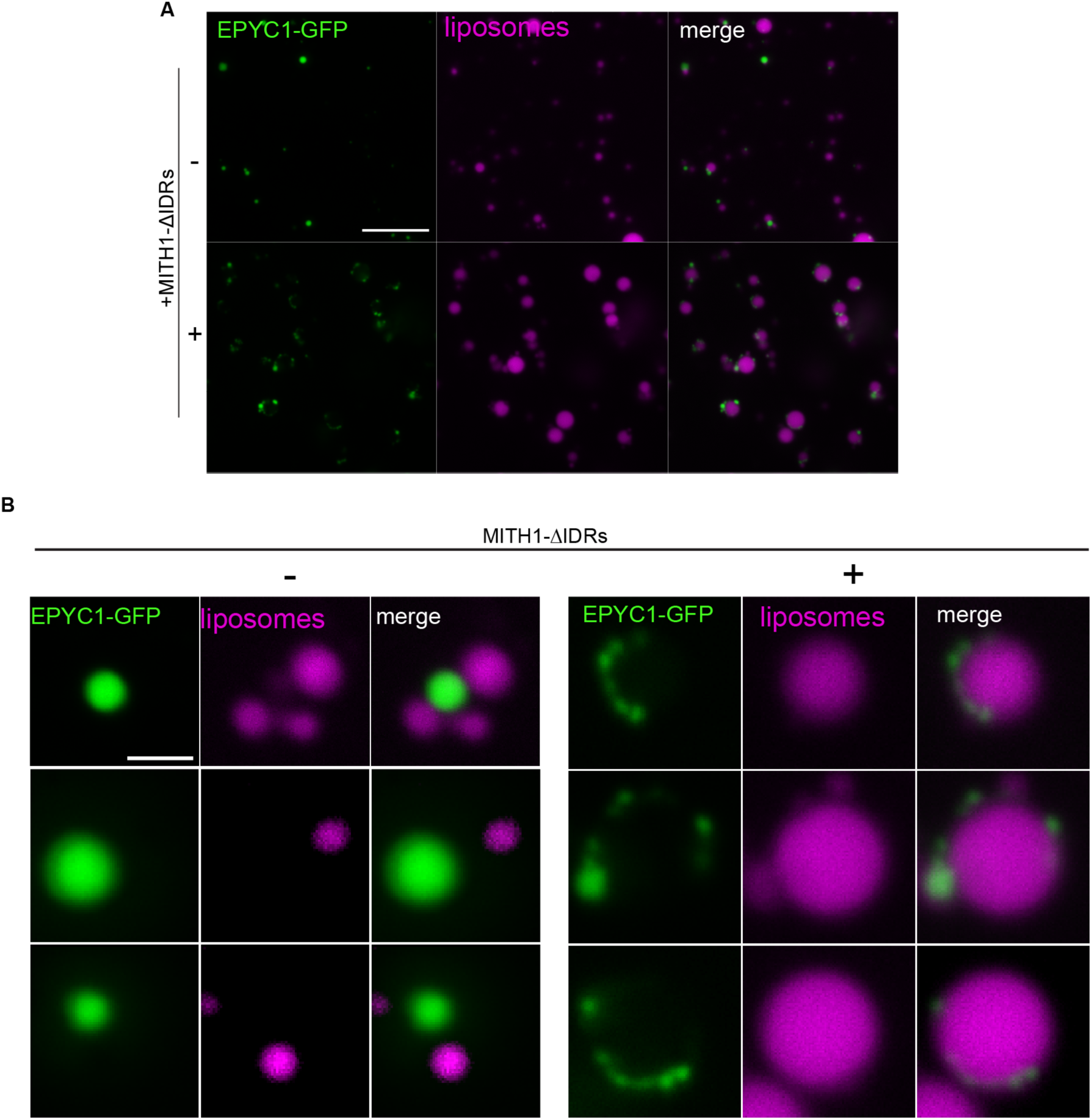
| MITH1 is sufficient to wet condensate onto membrane *in vitro.* **A** Full-field image of condensate and membrane with and without MITH1-ΔIDRs. Despite coming from the same preparation, liposomes are larger with the addition of MITH1. **B** Additional examples of condensate and membrane interaction with and without MITH1-ΔIDRs. Scale bar is 1 µm.

## Supplementary Tables

**Supplementary Table 1.** | Chlamydomonas strains used in this study.

| Chlamydomonas Resource Center ID | Strain Description | Mating Type | Source | Antibiotic Resistance |
| --- | --- | --- | --- | --- |
| CC-4533 | Wildtype | mt- | Previously published<sup>22</sup> | none |
| LMJ.RY0402.133670 | *mith1* | mt- | Previously published<sup>22</sup> | Paromomycin |
| CC-5938 | *mith1*;MITH1-Venus | mt- | Previously published<sup>22</sup> | Paromomycin, hygromycin |
| CC-6356 | *mith1*;MITH1*A7-Venus | mt- | Transformation of the *mith1* strain with a pRAM118 MITH1-Venus plasmid containing point mutations in the amphipathic helix region. | Paromomycin, hygromycin |
| CC-6357 | *mith1*;MITH1-CC-Venus | mt- | Transformation of the *mith1* strain with a pRAM118 MITH1-Venus plasmid containing deletions of the terminal IDRs and amphipathic helix region. | Paromomycin, hygromycin |
| CC-6358 | *mith1*;MITH1-ΔCC-Venus | mt- | Transformation of the *mith1* strain with a pRAM118 MITH1-Venus plasmid containing a deletion of the coiled-coil region. | Paromomycin, hygromycin |
| CC-6359 | *mith1*;MITH1-Δ1/2CC-Venus | mt- | Transformation of the *mith1* strain with a pRAM118 MITH1-Venus plasmid containing a deletion of half the coiled-coil region. | Paromomycin, hygromycin |

**Supplementary Table 2.** | Collection, refinement, and validation of cryo-EM model of Rubisco bound to MITH1.

| Data Collection |  |  |
| --- | --- | --- |
|  | Dataset #1 | Dataset #2 |
| Microscope | Thermo Fisher Titan Krios G3i |  |
| Camera | Falcon 4i |  |
| Energy Filter | Selectris X |  |
| Voltage (kV) | 300 |  |
| Spherical Aberration (mm) | 0 (Cs-corrected) |  |
| Magnification | 105 000 |  |
| Pixel size (Å) | 1.01 |  |
| Slit Width (eV) | 10 |  |
| Electron dose (e-/Å<sup>2</sup>) | 50 |  |
| Micrographs collected | 5,980 | 16,634 |
| Micrographs used | 5351 (89%) | 12981 (78%) |
| Initial particles picked | 229,617 | 1,165,316 |
| Particles after 2D classification | 91,670 | 705,788 |

| Final Refinement |  |
| --- | --- |
| Symmetry imposed | C1 |
| Final particle count | 101,807 |
| Global map resolution, masked (Å) | 2.91 |
| FSC threshold | 0.143 |

| Model Statistics |  |
| --- | --- |
| Composition (#) |  |
| Chains | 3 |
| Atoms | 9378 (Hydrogens: 4642) |
| Residues | Protein: 602 Nucleotide: 0 |
| Water | 0 |
| Ligands | 0 |
| Bonds (RMSD) |  |
| Length (Å) (# > 4σ) | 0.011 (0) |
| Angles (°) (# > 4σ) | 1.122 (0) |
| MolProbity score | 1.22 |
| Clash score | 1.81 |
| Ramachandran plot (%) |  |
| Outliers | 0.17 |
| Allowed | 3.89 |
| Favored | 95.95 |

| **Rama-Z (Ramachandran plot Z-score, RMSD)** |  |
| --- | --- |
| whole (N = 592) | -0.12 (0.34) |
| helix (N = 225) | 1.10 (0.35) |
| sheet (N = 85) | 0.07 (0.55) |
| loop (N = 282) | -1.09 (0.35) |
| **Rotamer outliers (%)** | 0 |
| **Cβ outliers (%)** | NA |
| **Peptide plane (%)** |  |
| Cis proline/general | 3.3/0.0 |
| Twisted proline/general | 0.0/0.0 |

## Methods

### Protein purification

MITH1-ΔIDRs and MITH1-CC were expressed in Sf9 insect cells and purified via N-terminal 6xHis-tag by GenScript.

MITH1-AH and EPYC1-GFP were purified in an *E. coli* expression system. Codon-optimized genes were cloned and inserted into the GST-tag-containing pGEX-6P-1 or pHUE vectors and transformed into BL-21 (DE3) competent cells. Next, 1 L cultures were induced at 0.8 OD with 1 mM IPTG and grown overnight at 16 °C. Cells were pelleted, resuspended in 50 mL TBS, pH 8.0 and lysed with two freeze-thaw cycles in liquid nitrogen. The lysate was sonicated for 5 minutes (3 seconds on, 6 seconds off, 60% power) and cleared by ultracentrifugation at 40,000 x g for 45 minutes. The supernatant was incubated for 1.5 hours with HisPur cobalt beads at 4 °C, washed 3x with 20 mM imidazole in TBS, and eluted with 300 mM imidazole. For GST-MITH1-AH, the GST tag was removed by PreScission protease used at 1 U per 100 µg protein overnight at 4°C and cleaned up by binding to anti-GST magnetic beads. Both proteins were further purified using Size Exclusion Chromatography on a Superdex column run on an Äkta Pure system.

Rubisco was purified from Chlamydomonas cells grown in Tris-Acetate-Phosphate (TAP) liquid media. Cells were grown to mid-log phase using the standard conditions described below and snap frozen in liquid nitrogen. Frozen samples were lysed by cryogenic grinding on a Cryomill (Retsch) at 20 oscillations per second for 20 minutes. Lysates were separated on a 10-30% sucrose gradient, collected with a piston gradient fractionator, and further purified by anion exchange chromatography on an Äkta Pure system. Rubisco was purified from *Arabidopsis* with the same method except for sample collection, in which whole leaves were snap-frozen and hand-ground with a mortar and pestle.

### Liposome generation and flotation assay

All lipids were purchased as powders from Avanti Polar Lipids. Liposomes were generated by mixing 5 mg/mL stocks of MGDG, DGDG, SQDG, and PG at a ratio of 5:3:1:1. Lipids were dried under a stream of nitrogen, left overnight in a vacuum chamber, and resuspended the following day in liposome buffer (10 mM HEPES, 50 mM KCl, pH 7.4) at a 3 mM concentration. The lipid mixture was then sonicated in a water bath for 20 minutes at room temperature. Then, purified protein at a concentration of 3 µM was added to liposomes to make a 1:1000 protein:lipid ratio. Equal volumes (100 µL) of protein lipid mixture and 60% sucrose solution in liposome buffer were combined to make a 30% sucrose proteoliposome layer in a low protein binding conical tube. Next, 200 µL of 25% sucrose in liposome buffer was gently layered on top of the 30% proteoliposome solution, followed by 50 µL of liposome buffer. The gradient was centrifuged at 18,000 x g for 20 minutes at 4 °C in a swinging bucket rotor. The liposome-containing fraction was clearly visible as a yellow band near the top of the tube. Bottom, middle, and top fractions were collected using a Hamilton glass syringe, and bottom (buffer), middle, and top (liposome) fractions were assessed for protein content using SDS-PAGE. Bands were quantified using ImageJ and enrichment in the top fraction was calculated as the total protein in the top fraction over the total protein in the top and bottom fractions combined. For the tubulation assay, 1 µM MITH1-AH or Venus control was added to liposomes, spotted on TEM grids and stained with negative stain as described below.

### Chlamydomonas culture conditions

Cells were maintained and cultured as previously described^23^. In brief, cells were grown to mid-log phase in TAP liquid media at air levels (0.04%) of CO_2_ and ∼200 µmol photons m^-2^ s^-1^. Liquid cultures were grown on an orbital shaker at 130 rpm at room temperature under continuous light. Sixteen hours prior to experiments, cultures were pelleted at 600 x g and resuspended in Tris-Phosphate (TP) liquid media at the same CO_2_ and light levels as the TAP cultures. Cultures were harvested for experiments at approximately 2 x 10^6^ cells/mL, as measured by a Countess Automated cell counter.

### Chlamydomonas transformation

To generate Chlamydomonas expressing truncated versions of MITH1, the previously described^19^ plasmid containing the MITH1 genomic sequence fused to Venus-3xFLAG in the vector backbone pRAM118 was modified with point mutations or deletions using site-directed mutagenesis or isothermal assembly. For transformation, *mith1* cells were grown to mid-log phase in TAP liquid media in air (0.04% CO_2_) using the standard conditions described above. The cells were centrifuged 5 minutes at 600 x g and washed twice with ice-cold MAX Efficiency transformation reagent (Invitrogen), then resuspended at 2 x 10^8^ cells/mL. For each transformation, 120 µL of cells were combined with 0.5-1 µg of plasmid linearized with restriction enzyme EcoRI (NEB) combined with 50 µg heat denatured carrier DNA (salmon sperm), incubated for 5 minutes on ice, and transferred to a pre-chilled electrocuvette (Bulldog Bio). Electroporation was carried out on a NEPA GENE 21 electroporator using the following settings: poring pulse 250, 8, 50, 2, 40, +; transfer pulse 20, 50, 50, 5, 40, +-. The cells were immediately transferred to 8 mL of TAP liquid media containing 40 mM sucrose, left rocking overnight at room temperature in dim light, and then plated on TAP agar plates containing selection antibiotic. After 5-7 days of growth in dim light, positive colonies were screened for Venus fluorescent signal on a typhoon scanner and verified by confocal microscopy (see below).

### Confocal microscopy

For all experiments, Chlamydomonas strains were cultured as described above. Cells were imaged live in 8-well imaging chambers (Ibidi) on a Nikon Ti2 Inverted Microscope with Yokogawa W1 and SoRa module. Movement of the cells was prevented by addition of a small block of TP-agar on top of the cells in each well. Cells were imaged with a 60x TIRF objective using excitation/emission wavelengths of 514/540 nm for Venus and 640/700 nm for Chlorophyll. Images were denoised and deconvolved using Nikon NIS Elements software.

### Transmission electron microscopy

For whole cell Transmission electron microscopy (TEM), strains were grown using the standard conditions described above at air levels of CO_2_, pelleted at 600 x g for 5 minutes, and fixed in 2.5% glutaraldehyde for 1 hour. Cells were then washed 3 x 5 minutes in MilliQ water, stained for 1 hour with a 1% OsO_4_ solution and again washed 3 x 5 minutes. Samples were serially dehydrated in ethanol solutions containing 50%, 75%, 95% and 100% ethanol followed by a 10-minute incubation in 100% acetonitrile. Dehydrated cells were left in a solution of 50% acetonitrile, 50% Quetol resin, then evaporated overnight. The samples were then embedded in epoxy resin containing 34% Quetol, 44% nonenyl succinic anhydride, 20% methyl-5-norbornene-2,3-dicarboxylic anhydride and 2% dimethylbenzylamine over 4 days of subsequent resin refreshes. On the final day, cells were pelleted at 18,000 x g for 20 minutes at 30 °C and cured at 60 °C for 48 hours. Resin blocks were cut with a Leica UCT Ultramicrotome and imaged on a Talos F200X S/TEM system.

For negative stain of *in vitro* proteins and liposomes, 5 µL of sample was placed on plasma cleaned TEM grids and stained with UranyLess negative stain. Grids were imaged on a Talos F200X S/TEM system.

### Spot test

Strains were grown to mid-log phase (2 x 10^6^ cells/mL) in TAP liquid media and pelleted at 600 x g for 5 minutes. Cells were resuspended at a concentration of 6 x 10^5^ cells/mL in TP liquid media and 10 µL was spotted onto TP agar plates. The plates were placed under constant light at 150 µmol photons m^-2^ s^-1^ for seven days in variable CO_2_ levels: very low CO_2_ at 40 ppm, air levels of CO_2_ at approximately 400 ppm, and high CO_2_ at 30,000 ppm.

### Sedimentation assay

Droplets were reconstituted *in vitro* in 20 µL reactions in 20 mM Tris-HCl, pH 8.0 and 50 mM NaCl and sedimented through centrifugation at 18,000 x g for 10 minutes. Proteins were used at the following concentrations: 2 µM Rubisco, 1 µM EPYC1, 1 µM MITH1-ΔIDRs, 1 µM MITH1-AH. The contents of the pellet and supernatant were analyzed by Coomassie-stained SDS-PAGE. The identical concentrations were used to generate the condensates used in the wetting assay.

### Peptide tiling array

Membranes displaying peptide arrays of 25 amino acid MITH1 peptides tiled every three amino acids were obtained from the MIT Biopolymer Core. Every fourth peptide was repeated, and the location of each peptide was randomized on the array. Membranes were activated by methanol and washed 3 x 10 minutes in Rubisco binding buffer (50 mM HEPES, 50 mM KOAc, 2 mM Mg(OAc)_2_, 1 mM CaCl_2_, 200 mM sorbitol, pH 6.8). The membrane was incubated overnight with 1 mg purified Rubisco labeled with NHS-488. To obtain labeled Rubisco, purified Rubisco was incubated with 3x molar excess NHS-488 for 1 hour at room temperature. The excess dye was removed by three passes through a resin-based dye removal column. After Rubisco binding, the membrane was washed 3 x 10 minutes in Rubisco binding buffer and imaged on an iBright imaging system. The images were quantified with ImageJ. The value of each repeated peptide sequence on the array was averaged.

### Fluorescence anisotropy

Peptides labeled with an N-terminal FITC were synthesized by GenScript. The anisotropy of 20 nM of peptides alone, peptides with 4 µM or 40 µM of Rubisco, and peptides with 40 µM BSA were measured on a Fluorolog-QM fluorimeter.

### Single particle cryo-electron microscopy

The Rubisco-MITH1 complex was prepared at respective concentrations of 0.5 mg/mL and 0.2 mg/mL in 10 mM HEPES, 150 mM KCl, 2 mM DTT, pH 7.4. UltraAUfoil R1.2/3 (Quantifoil Micro Tools GmbH) grids were glow-discharged for 30 seconds at 15 mA using a Pelco Easiglow. Then, 3 µL of the Rubisco-MITH1 complex sample was deposited on the front side of the grid. Excess sample was blotted for 4 seconds with blotting force 0, the chamber set at 12 °C and 95% humidity using a Vitrobot Mark IV (Thermo Fisher Scientific).

Data were collected on a Titan Krios G3i operating at 300 kV equipped with a Cs-corrector, Selectris X energy filter and Falcon 4i detector (Thermo Fisher Scientific). Data collection was set up on two different specimens using SmartScope^50,51^ with the following parameters: maximum beam image-shift radius of 4 µm with two exposures per hole, rolling target defocus of –0.8 to –2 µm and drift settling threshold at 1 Å/s. A total of 22,632 EER movies^52^ were collected at a 1.02 Å/pixel and a total dose of 50 e-/Å2 over a 3-second exposure with a 10 eV slit.

Data were processed using CryoSPARC^53^. Particles were picked using Topaz^54^. A few rounds of 2D classification were performed and only classes containing clear Rubisco were selected. For 3D refinement, the first reconstruction focused on Rubisco with D4 applied. Then, the symmetry was broken using symmetry expansion. A focus mask was created on the Rubisco-MITH1 interface and, after an initial local refinement, particles containing MITH1 were selected through 3D classification followed by additional local refinement. The 101,000 remaining particles containing the Rubisco-MITH1 complex were polished and subjected to a final local refinement. The final FSC resolution is estimated at 2.83 Å.

## Data and code availability

All data generated in this study are available within this article and its supplementary information files, with the exception of the cryo-electron microscopy map coordinates and model which are deposited in the Protein Data Bank (PDB) and the Electron Microscopy Data Bank (EMDB) (Accession #s: 10XV, EMD-75610).

## Acknowledgements

We acknowledge the support of the Confocal Imaging Facility, the Imaging and Analysis Center, and the Biophysics core with special thanks to Gary Laevsky and Sha Wang for help with confocal microscopy experiments, John Schreiber and Paul Shao for assistance with transmission electron microscopy and cryo-electron microscopy experiments and Venu Vandavasi for assistance with fluorescence anisotropy measurements. We thank the McCormick lab at Edinburgh for the gift of the Arabidopsis strains; Yana Kazachkova for providing the plant material; Clair A. Huffine of Insight Illustrations LLC for the scientific illustrations used in Figure 7A; Victoria Crans for assistance in generating figures and editing the manuscript; Marie Bao, as part of Life Science Editors, and Jackie Carozza and KC Farrell for editing the manuscript; Ned Wingreen, Trevor GrandPre, Silvia Ramundo, Aastha Garde, LaNell Williams and all of the Jonikas lab for helpful discussions and feedback. This work was supported by the Bill and Melinda Gates Foundation and United Kingdom Foreign, Commonwealth & Development Office grant INV-054558, National Science Foundation grant MCB-2410354, and by the Howard Hughes Medical Institute. This article is subject to HHMI’s Immediate Access to Research policy, which requires that this article be made publicly available as initial and revised preprints deposited on a designated preprint server under a CC BY 4.0 license.

## Author Contributions

S.L.E. and M.C.J. designed the study. S.L.E. and C.D. carried out experiments. J.B. carried out cryo-electron microscopy data collection and analysis. E.F. prepared samples for TEM and H.W. prepared and imaged samples for TEM. C.D.M. assisted with bioinformatic analysis. S.L.E. and M.C.J. wrote the manuscript with input from all authors.

